# Anti-CRISPR protein mediated degradation of Cas9 in human cells

**DOI:** 10.1101/2023.03.29.534776

**Authors:** Luisa Arake de Tacca, Joseph Bondy-Denomy, David Rabuka, Michael Schelle

## Abstract

Bacteriophages encode anti-CRISPR (Acr) proteins that inactivate CRISPR-Cas bacterial immune systems, allowing successful invasion, replication, and prophage integration. Acr proteins inhibit CRISPR-Cas systems using a wide variety of mechanisms. AcrIIA1 is encoded by numerous phages and plasmids, binds specifically to the Cas9 HNH domain, and was the first Acr discovered to inhibit SpyCas9. Here we report the observation of AcrIIA1-induced degradation of SpyCas9 and SauCas9 in human cell culture, the first example of Acr-induced degradation of CRISPR-Cas nucleases in human cells. Optimized expression of AcrIIA1 in human cells provided robust inhibition of SpyCas9 editing but had no impact on Type V CRIPSR-Cas12a, consistent with its binding site on the HNH domain. Targeted Cas9 protein degradation by AcrIIA1 could modulate Cas9 nuclease activity in human therapies. The small size and specificity of AcrIIA1 could be used in a CRISPR-Cas proteolysis-targeting chimera (PROTAC), providing a tool for developing safe and precise gene editing applications.

## Introduction

CRISPR (Clustered Regularly Interspaced Short Palindromic Repeats) arrays contain fragments of DNA that bacteria use as defense against invading nucleic acids (1,2). RNA-guided CRISPR-associated (Cas) nucleases identify invaders by first binding to a short protospacer adjacent motif (PAM) and then through Watson-Crick base-pairing, which leads to nucleic acid cleavage (3). Phages have developed CRISPR inhibitors that aid in evasion of the CRISPR defense and enhance the transmission of mobile genetic elements (MGE) (4). Anti-CRISPR (Acr) proteins inactivate the CRISPR-Cas immune system of bacteria (4–7). The first example of phage-encoded Acr proteins were found to inhibit the Class 1 Type I CRISPR-Cas systems (8,9). Shortly after this discovery, the first antagonists of Class 2 Type II CRISPR-Cas systems, including the clinically relevant SpyCas9, were identified in *Listeria* prophages (10,11). AcrIIA1 was revealed to be widespread across Firmicutes prophages and MGEs and has even been used as a marker for the discovery of new Acr proteins (10). AcrIIA1 is a broad-spectrum Cas9 inhibitor, capable of inhibiting multiple Cas9 orthologs (12). This broad inhibitory activity is due to the ability of AcrIIA1 to bind with high affinity to the conserved HNH domain of Cas9, specifically relying on the highly conserved catalytic residue H840. This allows AcrIIA1 to inhibit highly diverged Cas9 enzymes while other Acr proteins co-encoded with AcrIIA1 in *Listeria* phages, like AcrIIA4 and AcrIIA12 inhibit a much smaller range of Cas9 orthologs. AcrIIA1 inhibits multiple Type II-C Cas9 enzymes as well as the more common and therapeutically relevant Type II-A nucleases, including SauCas9 and SpyCas9 (12). Binding to a conserved domain and the resulting broad inhibition profile likely influenced the wide phylogenetic distribution of AcrIIA1 (10).

Other broad-spectrum Cas9 inhibitors, like AcrIIC1, also bind the conserved HNH domain (13). AcrIIC1 functions by trapping the DNA-bound Cas9 complex. Surprisingly, AcrIIA1 was found to stimulate degradation of catalytically active Cas9 protein in *Listeria* (12). In *Pseudomonas*, AcrIIA1 inhibited Cas9 without degrading the protein, suggesting that the degradation mechanism was specific to *Listeria*. While AcrIIA1 was able to weakly inhibit Cas9 in human cells, it was unable to degrade Cas9 *in vitro*, leading the authors to assume that the degradation mechanism relied on bacterial specific protein degradation machinery (12).

Here we demonstrate that AcrIIA1 induces Cas9 degradation in human cells. We show that AcrIIA1 degrades both SpyCas9 and SauCas9 but is unable to degrade or inhibit Type V CRISPR-Cas12a. To our knowledge, this is the first demonstration of Acr-induced Cas9 degradation in eukaryotic cells. This discovery allows for the development of therapeutic gene editing tools like CRISPR-Cas9 proteolysis-targeting chimera (PROTAC) (14,15). Acr-Cas9 PROTAC could be used to limit exposure of human genomes to Cas9 editing, reducing the potential for off-target effects and increasing the safety of gene editing therapies.

## Results and Discussion

### AcrIIA1 inhibits Cas9 gene editing in human cells

We transfected HEK293T human cells with a plasmid expressing SpyCas9 and a guide targeting the HBB locus and a second plasmid expressing AcrIIA1 (Fig 1A). Similar to previous results (10), AcrIIA1 encoded with native bacterial codons (AcrIIA1-bac) modestly inhibited SpyCas9 editing. However, expression of a human codon optimized version of the *acrIIA1* gene (AcrIIA1-hum) fully inhibited SpyCas9 editing. Editing at a known HBB off-target site (HBD) was also fully inhibited. Titration of AcrIIA1-bac plasmid showed a dose-dependent increase in SpyCas9 editing (Fig 1B). The AcrIIA1-hum construct was able to inhibit SpyCas9 editing at 0.5:1 ratio.

**Fig 1:**
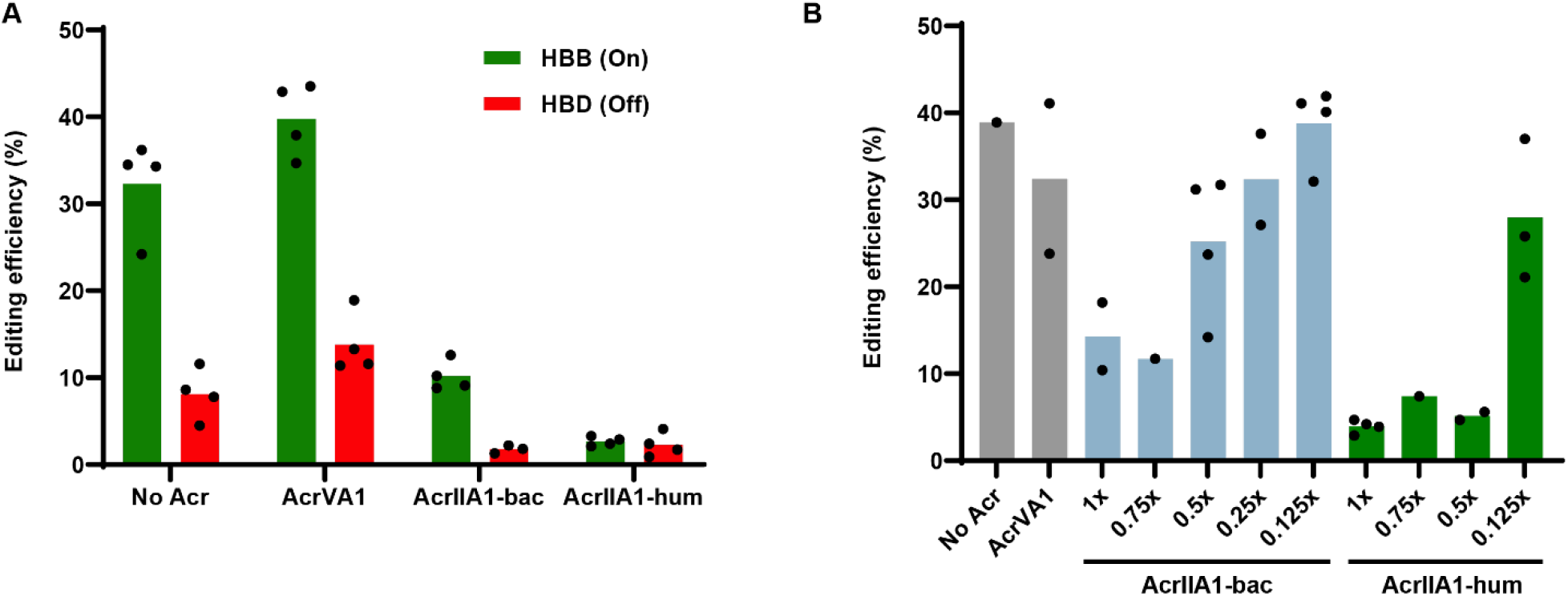
AcrIIA1 inhibits SpyCas9 editing in human cells. (A) Editing by SpyCas9 of the HBB gene and the closely related off-target site HBD. AcrIIA1-bac uses the native bacterial codons. AcrIIA1-hum is codon optimized for human expression. HEK293T cells were transiently transfected at a plasmid ratio of 1:2 SpyCas9:AcrIIA1 plasmid. (B) Dose dependent inhibition of SpyCas9 editing of HBB by AcrIIA1. “x” represents the fold plasmid amount of AcrIIA1 relative to SpyCas9. Total plasmid DNA transfected in each condition was constant.

### AcrIIA1 induces Cas9 degradation in human cells

We next sought to determine the mechanism of AcrIIA1 inhibition of Cas9 in human cells. We probed for the presence of SpyCas9 following expression in HEK293T cells using an anti-SpyCas9 monoclonal antibody. We expressed SpyCas9 and guide RNA alone or alongside various Acr constructs in HEK293T cells. Surprisingly, SpyCas9 was not detected when expressed with AcrIIA1 in human cells (Fig 2A). This is in contrast to co-expression of SpyCas9 with other Acr proteins including AcrIIA4, a strong SpyCas9 inhibitor (10), or AcrVA1, a Cas12a Acr that does not inhibit Cas9 (16). Neither AcrIIA4 nor AcrVA1 affected SpyCas9 expression in HEK293T cells. This result suggested that AcrIIA1 is stimulating the degradation of Cas9 in human cells, similar to the mechanism observed in *Listeria* (12). In contrast to AcrIIA1, AcrIIA4 is a potent SpyCas9 inhibitor that binds competitively to the PAM-interacting domain of SpyCas9 (17,18). The presence of SpyCas9 in the AcrIIA4 condition indicates that binding and inhibition of SpyCas9 is independent of degradation.

**Fig 2:**
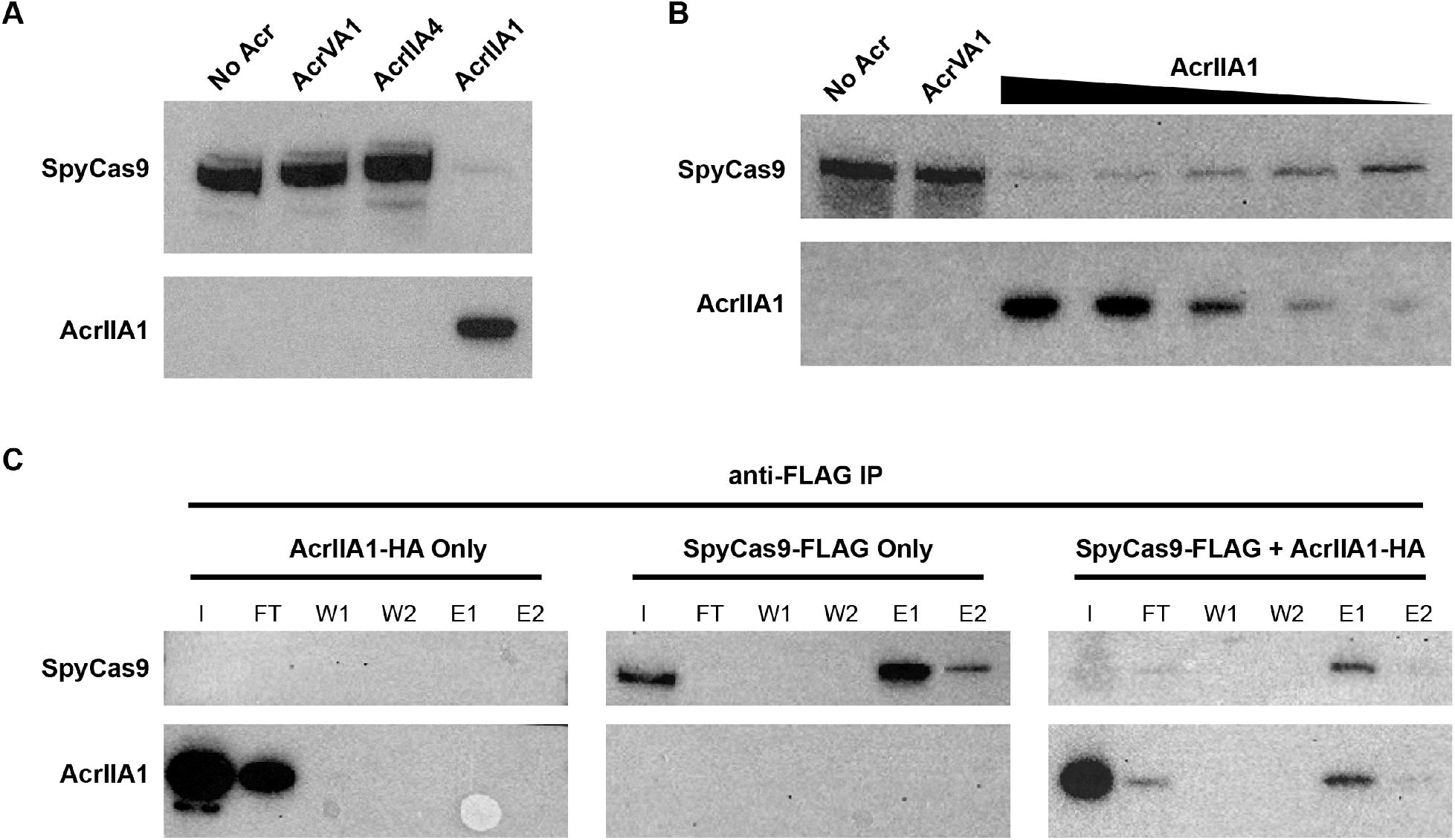
AcrIIA1-dependent degradation of SpyCas9. (A) Western blot showing AcrIIA1-dependent decrease in SpyCas9 protein level in HEK293T cells. Expression of AcrIIA4 or AcrVA1 do not show a decrease in SpyCas9 protein. (B) Western blot showing the dose-dependent decrease in SpyCas9 protein with increasing expression of AcrIIA1 in HEK293T cells. AcrIIA1 plasmid is dosed from 1x to 0.125x relative to SpyCas9 plasmid. (C) Western blots of anti-FLAG immunoprecipitations pulling down FLAG-tagged SpyCas9 and probing for SpyCas9 and AcrIIA1. AcrIIA1-HA alone is fully eluted in the flow through (FT). SpyCas9-FLAG efficiently binds to the anti-FLAG beads and is eluted (E1 and E2). Co-expression of SpyCas9-FLAG and AcrIIA1-HA (1:0.125 plasmid ratio) results in lower SpyCas9. AcrIIA1 binds and elutes (E1 and E2) along with the residual SpyCas9. I = input, FT = flow through, W1 = wash 1, W2 = wash 2, E1 = elution 1, E2 = elution 2.

We next assessed the dose-dependence of AcrIIA1-induced Cas9 degradation. Plasmid encoding AcrIIA1 tagged with an HA epitope (S1 Fig) was titrated and transfected into HEK293T cells along with a plasmid expressing SpyCas9 and guide RNA (Fig 2B). An anti-HA antibody shows an increase in AcrIIA1 expression with increasing plasmid concentration. SpyCas9 protein concentration is inversely correlated with AcrIIA1 expression, consistent with AcrIIA1-incuded degradation. The SpyCas9 protein concentration also correlates with the editing observed in Fig 1B, with increased editing and SpyCas9 protein at 0.125-fold AcrIIA1 plasmid concentration.

### AcrIIA1 binds SpyCas9 in human cells

To determine if AcrIIA1 is directly binding SpyCas9 in human cells, we co-immunoprecipitated AcrIIA1 and SpyCas9 (Fig 2C). Lysates from HEK293T cells transfected with plasmids encoding HA-tagged AcrIIA1 and FLAG-tagged SpyCas9 were immunoprecipitated with magnetic beads conjugated to an anti-FLAG antibody to pull down the SpyCas9 protein. HEK293T cells were transfected with SpyCas9-FLAG and AcrIIA1-HA plasmids at either a 1:1 ratio (S2 Fig) or 1:0.125 ratio (Fig 2C). SpyCas9 is barely detectable in the lysate at both AcrIIA1 ratios, though immunoprecipitation enriched for remaining Cas9. In both conditions, AcrIIA1 co-elutes with SpyCas9, indicating direct binding between SpyCas9 and AcrIIA1 in human cells. To assess if AcrIIA1-induced SpyCas9 degradation leads to truncation products, we probed a Western blot using an anti-SpyCas9 antibody. We did not observe any obvious degradation products when AcrIIA1 was added, only a decrease in overall protein levels (S3 Fig).

### AcrIIA1 induces degradation of Cas9 orthologs in human cells

Given its wide inhibition spectrum observed in bacteria, we tested AcrIIA1 for inhibition of SauCas9 in human cells. In bacteria, SauCas9 is inhibited to a lesser degree than SpyCas9 by AcrIIA1 (12). Similarly, we observed that AcrIIA1 is only able to modestly inhibit SauCas9 in human cells (Fig 3A). This is in contrast to the total inhibition seen with SpyCas9 (Fig 1A). Like with SpyCas9, the inhibition is dose-dependent, with a lower concentration of AcrIIA1 plasmid resulting in less inhibition of SauCas9 editing. Despite the modest inhibition of SauCas9 editing by AcrIIA1, the Acr still efficiently induces degradation of SauCas9 protein (Fig 3B). Indeed, at the 1:1 plasmid ratio, SauCas9 is barely detectable in HEK293T cell lysates. Unlike with SpyCas9, SauCas9 protein levels are fully restored at the 1:0.125 plasmid ratio, indicating that the AcrIIA1-induced degradation of SauCas9 is weaker than with SpyCas9. These results indicate that even highly diverged Cas9 orthologue are susceptible to the degradation mechanism employed by the AcrIIA1 family. Given the diversity of the AcrIIA1 family, we speculate that an AcrIIA1 homologs exist that would provide more robust inhibition and degradation of SauCas9.

**Fig 3:**
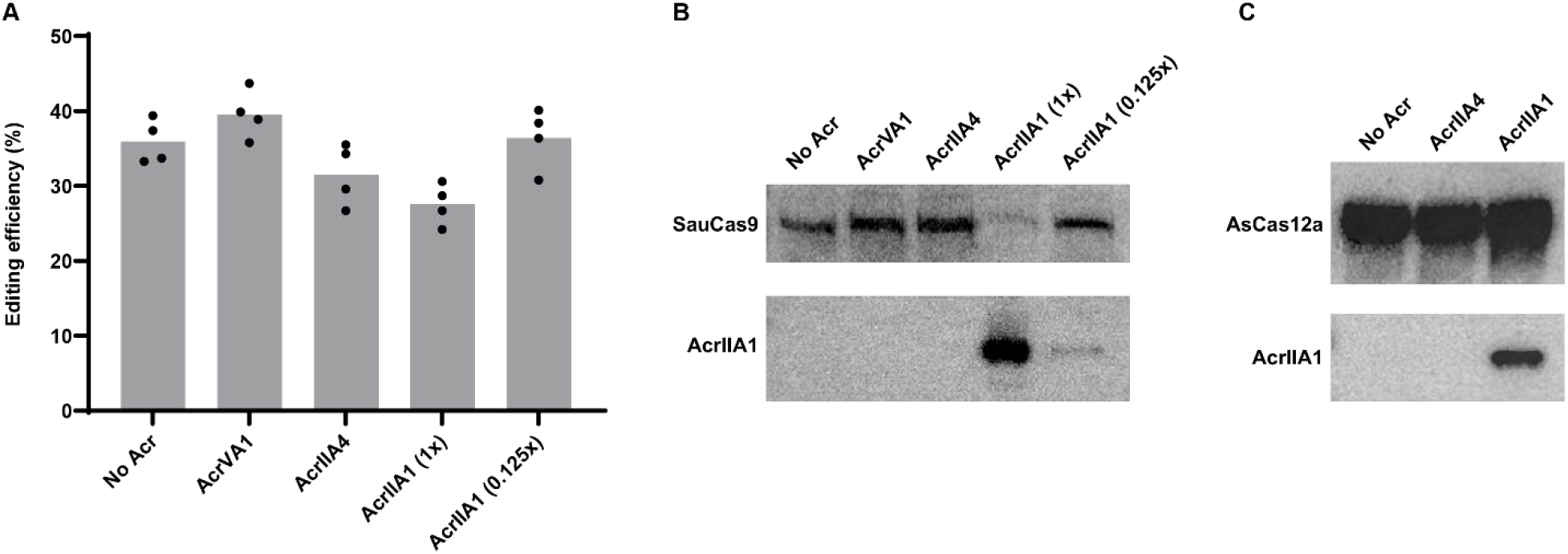
AcrIIA1 induces degradation of SauCas9 but not Cas12a. (A) Editing efficiencies for SauCas9 targeting Chrm2. AcrIIA1 only modestly inhibits SauCas9 at a 1:1 plasmid ratio (AcrIIA1 (1x)). (B) Western blot of SauCas9 co-expressed with various Acr proteins. Co-expression of AcrIIA1 induces degradation of SauCas9 similarly to SpyCas9. AcrIIA4 and AcrVA1 do not affect SauCas9 protein concentrations. (C) Western blot of AsCas12a co-expressed with various Acr proteins. AsCas12a protein concentration is not affected by either AcrIIA1 or AcrIIA4.

In bacteria, AcrIIA1 was unable to inhibit CRISPR-Cas systems beyond the Type II Cas9 family. We tested if AcrIIA1 was able to degrade the Type V nuclease AsCas12a, which lacks an HNH domain and the catalytic residue AcrIIA1 is known to interact with (12). Co-expression of AsCas12a and AcrIIA1 from plasmids at a 1:1 ratio shows no degradation of AsCas12a (Fig 3C). Probing for AcrIIA1 shows that the protein is expressed, indicating that there is no interaction between the Type V nuclease and AcrIIA1. Taken together, these results indicate that AcrIIA1 broadly inhibits and induces the degradation of Cas9 nucleases in human cells and that this mechanism is specific to Type II CRISPR-Cas systems.

## Discussion

In this work, we show for the first time that an anti-CRISPR protein is capable of inducing the degradation of a CRISPR-Cas nuclease in human cells. Destabilization or degradation of a Cas protein by an Acr is an uncommon mechanism. AcrIIA1 was previously shown to inhibit and induce degradation of Cas9 orthologs in *Listeria* (12). Key binding residues were elucidated on both the Acr and Cas, explaining the broad phylogenetic distribution of the AcrIIA1 family and breadth of Cas9 inhibition. While the exact mechanism of AcrIIA1-induced Cas9 degradation remains unknown, the authors concluded that the degradation mechanism was likely to be limited to certain bacterial species where Cas9 and AcrIIA1 are naturally found.

In this report, we show that AcrIIA1 induces degradation of SpyCas9 and SauCas9 by direct binding in human cells. This surprising observation could be used to develop a Cas9 PROTAC, which is capable of controlled Cas9 degradation, similar to previously engineered auxin inducible degron fusions (15). Altogether, the ability of a single protein domain (∼80 amino acid C-terminal domain of AcrIIA1) to inhibit and degrade numerous Cas9 proteins in human cells suggests that this protein is either a protease, a Cas9 destabilizer, or interfaces surprisingly well with human protein degradation machinery. AcrIIA1 binds tightly to the Cas9(D10A) nickase (12,19,20), commonly used in base editing applications (21), suggesting that this gene editing tool could also be degraded. The utility of irreversibly degrading (as opposed to inhibiting) Cas9-based tools could provide a robust stand alone “Cas9 off-switch” or be paired with strong inhibitors of DNA-binding (e.g. AcrIIA4), analogous to the approach used by bacteriophages (22,23).

## Materials and Methods

### Plasmid Cloning

Three plasmids containing AcrIIA1 were constructed expressing either: a native bacterial codon *acrIIA1* (AcrIIA1-bac), a human codon-optimized *acrIIA1* (AcrIIA1-hum), or AcrIIA1-hum with an HA-tag on the N-terminus (AcrIIA1-HA). These were ordered as gene fragments from Twist Bioscience and cloned into Twist’s CMV expression vector using HindIII and BamHI restriction sites. AcrVA1 and AcrIIA4 used as controls were codon-optimized for human expression and ordered and cloned exactly as AcrIIA1.

The SpyCas9 plasmid was purchased from Genscript with BbsI cloning sites for guide addition. An HBB guide was added to the SpyCas9 plasmid through the oligo anneal protocol provided from Dr. Feng Zhang’s lab available online under “PX330 cloning protocol”. The oligos used to make the HBB target are listed as Supporting information.

The SauCas9 plasmid was purchased from Genscript with BsaI cloning sites for guide addition and the sequence is the same as PX601 from Dr. Feng Zhang’s lab. Guide cloning protocol for the Chrm2 also follows “PX330 cloning protocol” and uses oligo anneal as the methods. The oligos used for Chrm2 guide listed as Supporting information.

### Sequencing

For sequencing of the endogenous regions assessed for editing efficiencies, we used primers that annealed to each specific region. The off-target region shown in Fig 1 is located in the HBD locus and the sequence assessed is located at the Intergenic Position: chr2:116069276-116069298:+ and the sequence is G**G**GAACGTGGATGAAG**C**TGG (*AGG)* in which the bold letters represent mismatches to the guide. Each region was amplified using the primers listed in the Supporting information and checked on a 2% agarose gel for purity. They were cleaned up using a PCR clean up kit from Zymo (CAT D4033) and submitted to sanger sequencing using the sequencing primer provided above. TIDE analysis was performed following the published method (24) and performed according to recommendations. All PCR and sequencing primers are listed in the Supporting information.

### HEK293T transfection

Cas9 and guide plasmids and the Acr plasmids were tested for activity in HEK293T cells following plasmid transfection using Mirus Transit X2 reagent. Tests were performed in 96 well plates transfected with 100 ng of nuclease expression vector and varying amounts of Acr vectors depending on the experiment following the Mirus Transit X2 transfection recommendations. Samples were incubated for 72h and harvested with Quick Extract (Lucigen). Genomic DNA was amplified and sequenced as described above.

### Western Blot and Immunoprecipitations

We used NP40 lysis buffer (50mM Hepes Ph 7.5, 150 mM KCl, 2mM EDTA, 0.5% NP40 and 1mM DTT). Before use, we add 1 Roche complete tablet for 10mL of buffer. Samples were loaded using SDS loading buffer (100mM Tris-Cl pH 6.8, 4% SDS, 0.2% bromophenol blue, 20% glycerol, 200mM of DTT for 10mL of water). Transfected HEK293T cells are lysed, and we performed Bradford to normalize gel loading amounts. We used the iBind system to transfer the gel before blotting and iBlot for blotting the western blots. For SpyCas9 we used a mouse monoclonal antibody from Cell Signaling (CAT 65832) at 1:1000 amount. For SauCas9, we used a rabbit polyclonal antibody from Millipore Sigma (CAT AB356480) at 1:1000 dilution. For HA-tagged AcrIIA1-HA, we used a rabbit monoclonal anti-HA antibody from R&D systems (CAT MAB0601).

For the anti-FLAG IP, we used anti-FLAG M2 Affinity Gel from Merck (CAT A2220) following manufacturer’s instructions. For HSP90 we used a polyclonal antibody raised in rabbit from Cell Signaling (CAT 4874) at a 1:1000 dilution.

## Acknowledgements

We want to thank the members of Acrigen Biosciences for their assistance and helpful discussion.

## Anti-CRISPR protein mediated degradation of Cas9 in human cells

Luisa Arake de Tacca^1^, Joseph Bondy-Denomy^2^, David Rabuka^1^, Michael Schelle^1*^

## Supporting Information

### Oligos and primers used in the study

HBB guide FWD: CACCGGTGAACGTGGATGAAGTTGG

HBB guide REV: AAACTGTGGGGCAAGGTGAACGTGC

Chrm2 guide FWD: CACCGGAGGTCTGGCAGCCAAGATGG

Chrm2 gudie REV: AAACCCATCTTGGCTGCCAGACCTC

HBB Amp FWD: CGATCCTGAGACTTCCACACTG

HBB Amp REV: CCAATCTACTCCCAGGAGCAGG

HBB Seq: CCAATAGGCAGAGAGAGTCAG

Chrm2 Amp FWD: GCGAATGCTGCTGTCTGATCAT

Chrm2 Amp REV: AGGAAGCCAGAGGTAATTCAGGT

Chrm2 Seq: AGGAAGCCAGAGGTAATTCAGGT

OT2 Amp FWD: GCAGTCCTGTGTTGAGAATA

OT2 Amp REV: ACCGTTTGTCTTTCTGTACC

OT2 Seq: CATCAATCCCATTATTACAC

**S1 Fig.**
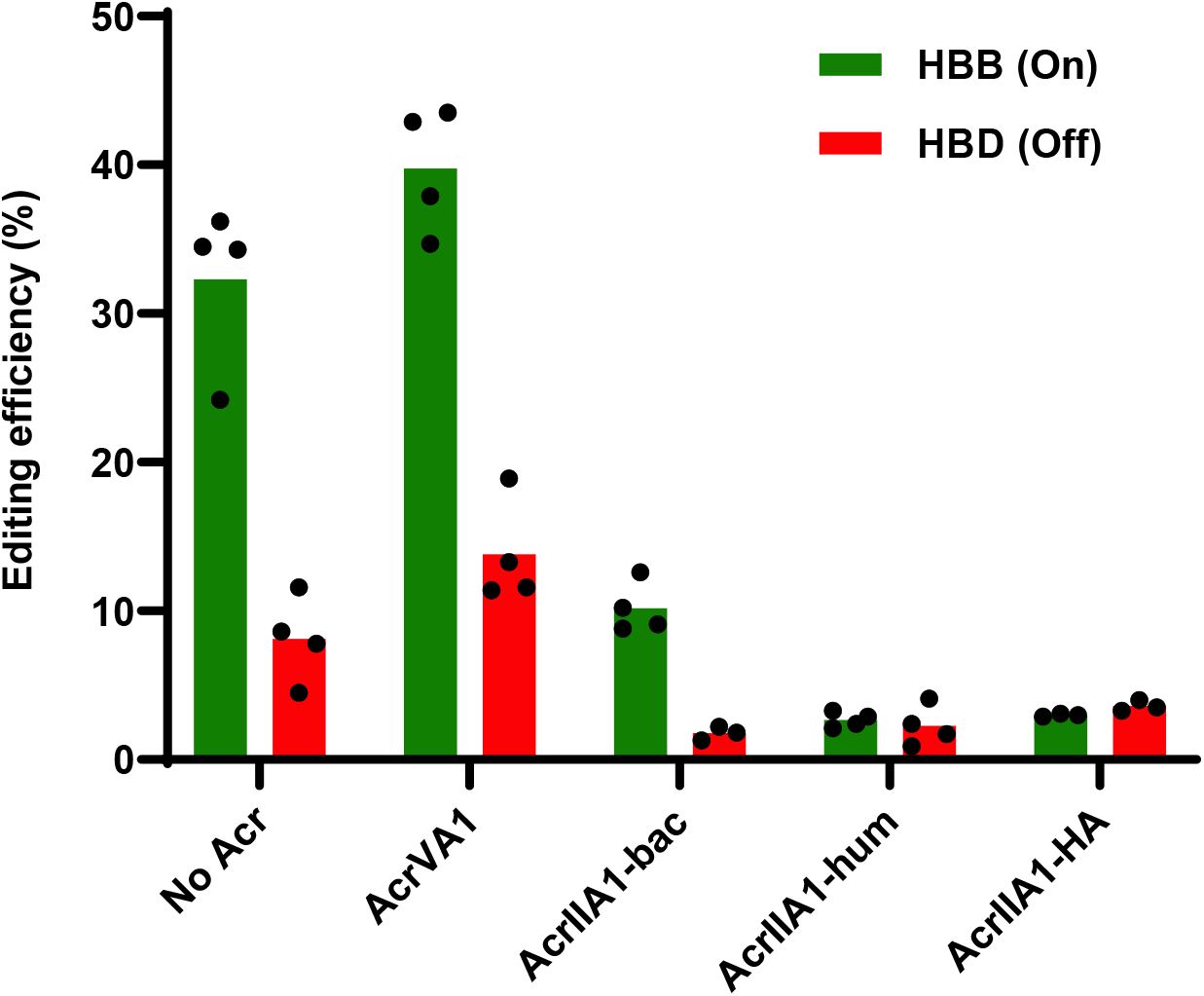
AcrIIA1-HA inhibits similarly to AcrIIA1-hum. Editing by SpyCas9 of the HBB gene and the closely related off-target site HBD. AcrIIA1-bac uses the native bacterial codons. AcrIIA1-hum is codon optimized for human expression. AcrIIA1-HA is the AcrIIA1-hum with an HA tag. HEK293T cells were transiently transfected at a plasmid ratio of 1:2 SpyCas9:AcrIIA1 plasmid.

**S2 Fig.**
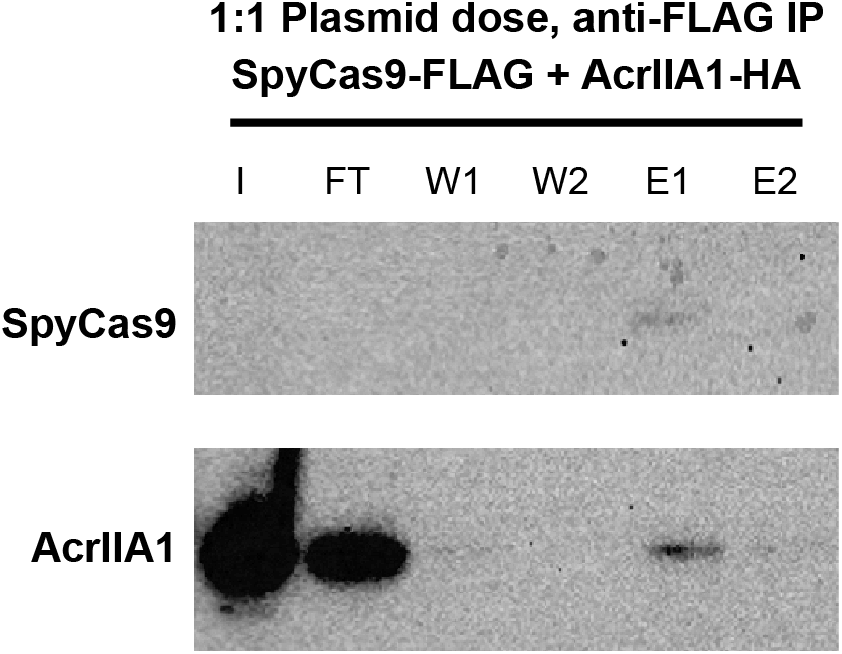
AcrIIA1 binds SpyCas9. Western blot of anti-FLAG immunoprecipitations pulling down FLAG-tagged SpyCas9 and probing for SpyCas9 and AcrIIA1. Co-expression of SpyCas9-FLAG and AcrIIA1-HA (1:1 plasmid ratio). AcrIIA1 binds and elutes (E1) along with the residual SpyCas9. I = input, FT = flow through, W1 = wash 1, W2 = wash 2, E1 = elution 1, E2 = elution 2.

**S3 Fig.**
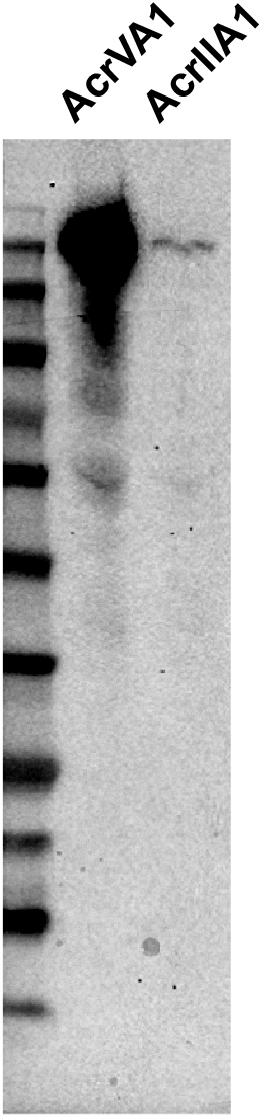
No SpyCas9 degradation products detected. Western blot showing AcrIIA1-dependent decrease in SpyCas9 protein level in HEK293T cell lysates compared to AcrVA1. No degradation products are seen in the AcrIIA1 condition that are not present in the AcrVA1 lysate. SpyCas9 is detected using a monoclonal anti-SpyCas9 antibody.

